# Organoid-derived adult human colonic epithelium responds to co-culture with a probiotic strain of *Bifidobacterium longum*

**DOI:** 10.1101/2020.07.16.207852

**Authors:** Emma Lauder, Kwi Kim, Thomas Mitchell Schmidt, Jonathan Louis Golob

## Abstract

In germ-free animals the lack of microbes within the gut results in a propensity to mucosal inflammation among other immune deficits. This suggests microbes are essential for the healthy function of the human gut, but we have lacked a reproducible mechanistic model of interactions between human colonic epithelium and the anaerobic microbes within the gut. To establish the physiological effect of a common anaerobe in the human gut, we co-cultured a probiotic strain of *Bifidobacterium longum* (35624) with organoid-derived adult human colonic epithelium in asymmetric gas conditions (anoxic apically, 5% oxygen basolaterally) and compared to axenic (‘germ-free’) epithelium. Bacteria proliferated and retained their normal cellular morphology in the presence of the human colonic epithelium. The human colonic mucosa retained trans-epithelial electrical resistance (TEER) consistent with an intact epithelium but lower than in axenic conditions. Changes in TEER corresponded to changes in Claudin-family gene expression. Inflammation was repressed in co-culture as compared to axenic, with reduced expression of executor and pyroptosis caspases; reduced expression of activators and increased expression of inhibitors of *NFKB*; reduced expression of toll-like-receptors (*TLR*s) and increased expression of *TOLLIP* (a negative regulator of TLRs). Consistent with the presence of actively fermenting bacteria that produce lactate and acetate but do not produce butyrate, *PPARA* expression was increased while *PPARG* expression was reduced. As in germ-free animal experiments, axenic human colonic mucosa is poised for inflammation. Co-culture with *Bifidobacterium longum* resolved the pro-inflammatory state while modulating barrier function (via Claudin genes) and cellular energetics (via *PPARA* and *PPARG* genes).

**Importance:** Experiments in animals demonstrate the importance of microbes with the gut for the health of the gut. Many of the microbes within the healthy gut are strictly anaerobic and cannot grow in the presence of oxygen; human tissues require some oxygen to live. Here we observe the effect of growing an anaerobic bacterium - *Bifidobacterium longum* - typically found in the human gut with human colonic tissue. We observed that the human tissue responded favorably to being cultivated with the microbe as compared to being alone. This experiment confirms human colonic tissues reduce their inflammation in the presence of a bacteria typically found in the gut.

## Observation

Animals without a gut microbiome are known to have a dysfunctional gut, including lowered GI motility (1), altered ionic transport across the mucosal barrier (2), chronic diarrhea (3), and difficulty absorbing nutrients (3). Germ-free animals also fail to properly develop innate or adaptive immunity (4, 5), have impaired reproductive fitness (6), and deficiencies in bone and neural development (7, 8). These animal-model studies are the basis for our belief that the presence of a gut microbiome is critical for human health. However we have previously lacked an adequate *human* model of host-microbiome interactions to verify this belief.

Organoid-derived epithelia allow for reproducible and mechanistic investigation of interactions between anaerobic bacteria and human colonic epithelium. Organoid-derived human colonic epithelia retain the genetic and epigenetic state of the tissue from which it was derived (9). This model was recently extended to allow for co-culture of the human epithelium with anaerobic microbes typically found within the human gut (10). To establish the physiological effect of a common anaerobe in the human gut, we co-cultured a probiotic strain of *Bifidobacterium longum* (35624) with organoid-derived adult human colonic epithelium in asymmetric gas conditions (anoxic apically, 5% oxygen basolaterally) and compared its behavior to axenic (‘germ-free’) epithelium.

In asymmetric oxygen conditions, both the microbes and organoid-derived human epithelium remained metabolically active. After 24 hours, an inoculum of 10^6^ colony forming units (CFU) per microliter of *B. longum* proliferated to a median of 1.9×10^8^ CFU/mL (Fig. 1A) in co-culture with human colonic epithelium, similar to microbes in traditional anaerobic culture conditions. The epithelium retained trans-epithelial electrical resistance at a level consistent with an intact monolayer, but at a reduced level as compared to axenic conditions (Fig. 1B).

**Figure 1:**
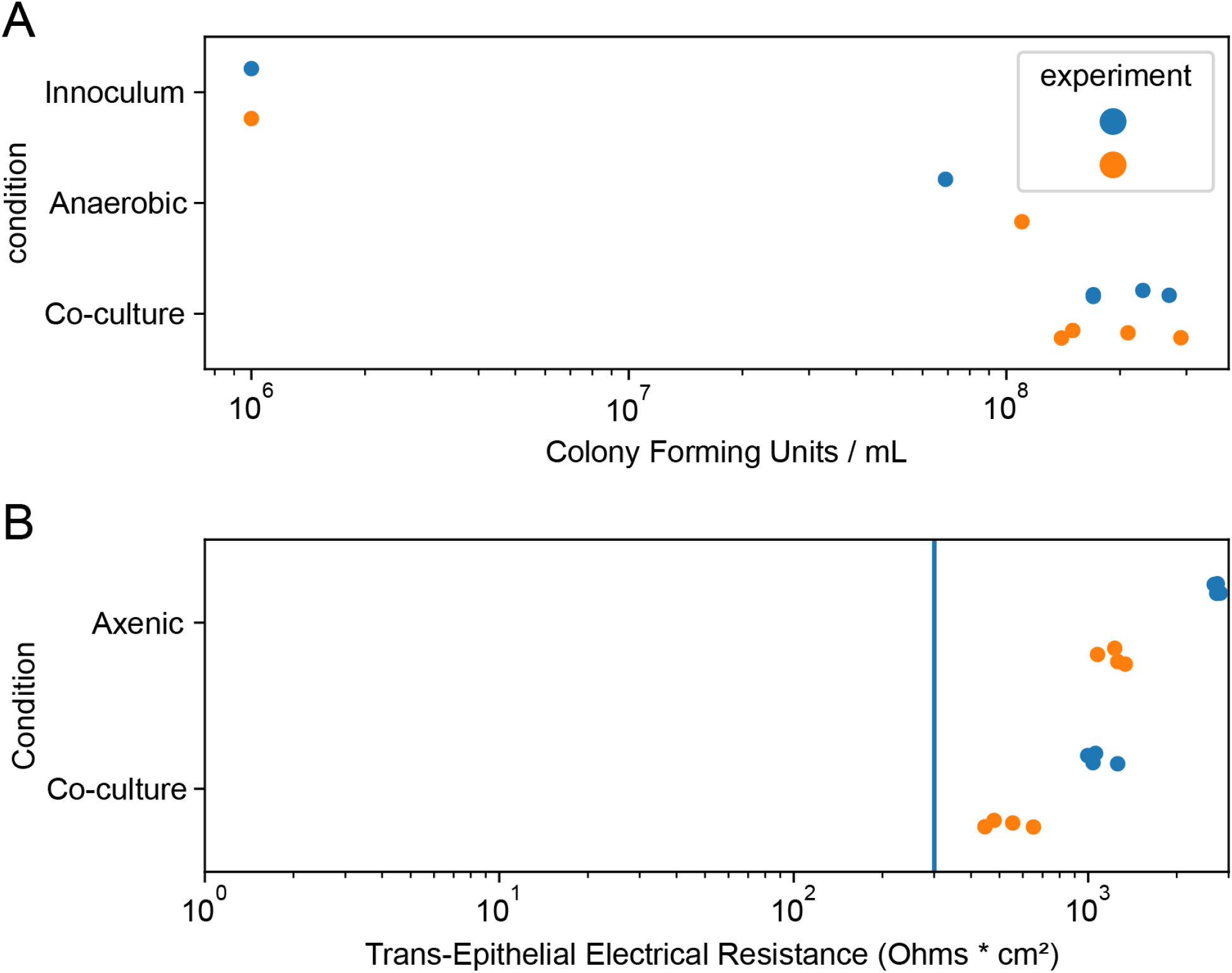
**The number of** microbes (A) and electrical resistance of human epithelium (B) after 24 hours of co-culture in asymmetric gas conditions, as compared to axenic controls.

Co-culture (as compared to microbe-free axenic culture) resulted in multiple changes in gene expression consistent with reduced inflammation (Fig. 2). Compared to axenic conditions, co-culture with *Bifidobacterium longum* (35624) resulted in reduced expression of executor and pyroptosis caspases (Fig. S1). NFκB is a master-regulator transcription factor of cellular inflammation (11). NFκB activators were downregulated and NFκB inhibitors were upregulated with co-culture (Fig. S2). Toll-like receptors (TLRs) recognize microbial molecular patterns (12); inflammatory TLRs were downregulated and TOLLIP (a negative negative regulators of TLRs) was upregulated (Fig. S3).

**Figure 2:**
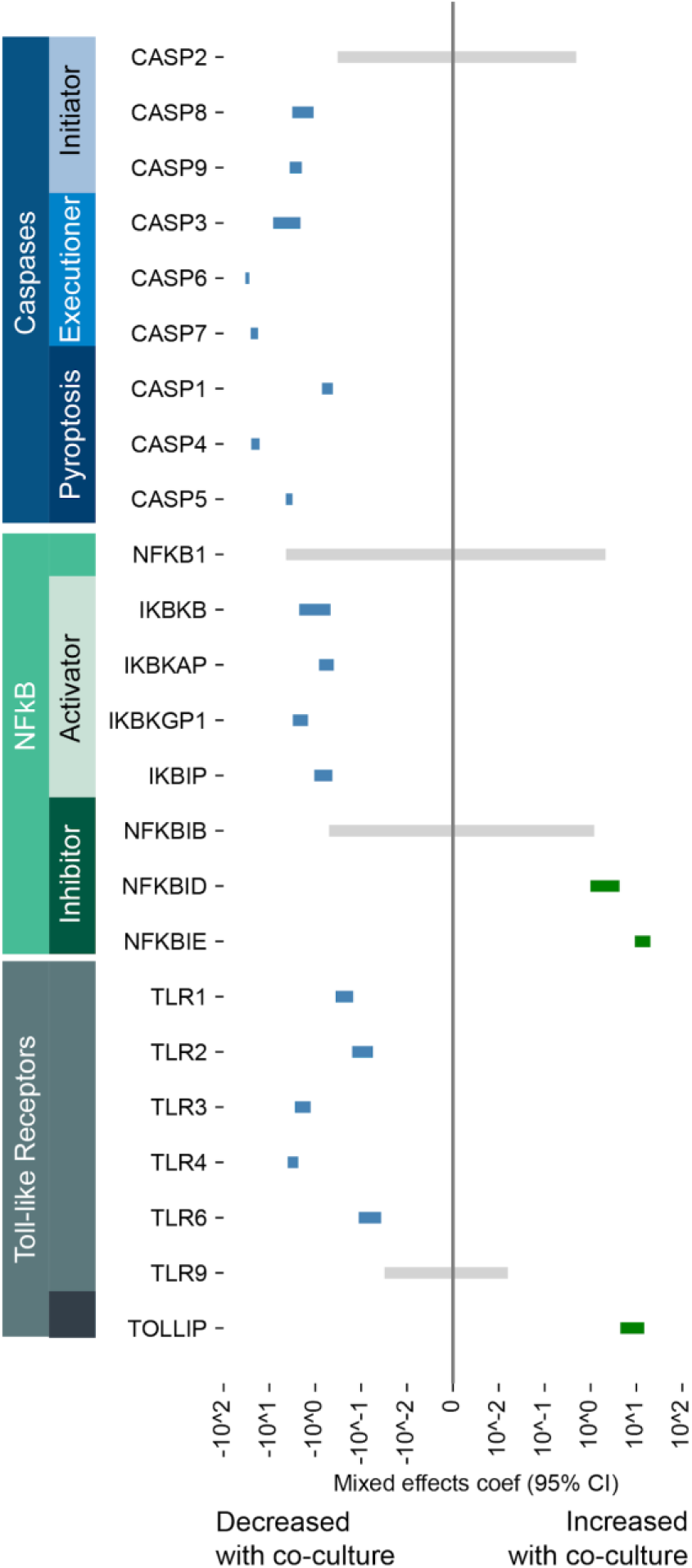
Changes in regulators of inflammation and apoptosis with co-culture. The 95% confidence interval for the linear-mixed / random effects model coefficient is depicted for co-culture as the independent variable, and fragments per kilobase of transcript length and megabase of sequence for each transcript. If the confidence interval does not cross zero, expression was significantly reduced (blue) or increased (green) when comparing co-culture to axenic control conditions.

Co-culture affected the expression of genes associated with the tight junctions, which corresponded to the observed change in TEER. Claudin-family gene expression changed with *CLDN23*, *CLDN1*, and *CLDN2* downregulated and *CLDND1* and *CLDND2* upregulated with co-culture as compared to axenic (Fig. S4).

Cellular energetics were also affected by co-culture. The peroxisome proliferator activated receptor (PPAR) family of transcription factors are master regulators of cellular respiration. Consistent with co-culture with a bacterium that produces lactate and acetate as fermentation products, *PPAR-γ* expression was downregulated and *PPAR-α* expression was upregulated as compared to axenic conditions (Fig. S5).

### Here we observe co-culture with *Bifidobacterium longum* resolved the pro-inflammatory state of axenic epithelium while modulating barrier function (via Claudin genes) and cellular energetics (via *PPAR* genes)

Our observations with organoid-derived human colonic epithelium *in-vitro* are consistent with previous findings in germ-free animal models (1–8). We also observed that both the anaerobic microbes and organoid-derived human colonic epithelium survived and remained metabolically active when co-cultured together in asymmetric oxygen conditions.

Our observation demonstrates the promise of this *in vitro* systems biology experimental approach. We feel organoid-derived human colonic epithelium is more predictive of true physiologic responses than immortalized colonic carcinoma cell lines. Compared to animal models, this approach is more efficient in time and expense and does not require extrapolation from the model organism to human biology. Even with the introduction of a single microbial species typically found in the human gut, we were able to observe changes in the transcriptional state of the colonic epithelium consistent with prior *in vivo* model organism experiments.

Compared to the host-microbe interactions in a healthy adult human colon, this experiment was quite simplified with only one microbial strain present. Further, this strain contains NADH-oxidase and superoxide dismutase (13) and therefore may be more micro-aerotolerant compared to other anaerobes found in the human gut. There is no obvious barrier to future experiments with more complex microbial communities (either adoptively transferred from the lumen of the human gut, or constructed communities in culture), but future experiments will have to determine the phylogenetic and physiological range of anaerobic microbes that survive and remain metabolically active in this co-culture system.

In this observation, we made extensive use of RNA-sequencing to determine the transcriptional state of the human colonic epithelium. This systems biology approach is suitable to more definitive measures of the state of the human epithelium, including immunochemistry, colorimetric assays of transcriptional factor and enzymatic activity, and Western blot.

The combination of organoid-derived human colonic epithelium with asymmetric oxygen culture conditions is a promising novel model system for future human host-microbe systems biology experiments.

## Methods

### Bacterial preparation

Bifidobacteria Specific Media (BSM) broth (Sigma-Aldrich; 90273) was inoculated with *Bifidobacterium longum* strain 35624 and cultured for 18 hours while shaking at 37C. The bacterial suspension was washed and diluted in DMEM/F-12, GlutaMAX supplement (Thermofisher Catalog# 10565018).

### Human Colonic Epithelium from Organoids

Colonic stem cells (University of Michigan TTML colonoid line 87) were plated onto collagen-coated transwells in accordance with the protocol from the Translational Tissue Modeling Laboratory (http://www.jasonspencelab.com/ttml). The stem cells were differentiated, as per (14). Differentiation was completed in a 5% oxygen 5% CO2 balance N2 environment to acclimate to physiological oxygen conditions. Experimental conditions were added 18 hours after first TEER measurement if TEER was determined to be present (TEER>330Ohms/cm2) indicating an intact monolayer of cells.

### Culture Conditions

Differentiation Media (14) was removed from the apical side of the transwell insert and either DMEM/F-12, GlutaMAX supplement (Thermofisher Catalog# 10565018) (axenic) or an inoculum of *Bifidobacterium* of a known starting OD were added. Co-culture occurred at 37 C. The basal compartment was maintained with a 5% oxygen, 5% CO2 balance N2 environment. The apical chamber was maintained in anaerobic conditions in a 5% CO2 balance nitrogen environment.

### Trans-epithelial Electrical Resistance (TEER)

TEER was recorded using the electrical Volt/Ohm meter (World Precision Instrument).

### Gene expression

After 24 hours of either axenic growth or co-culture with Bifidobacterium, RNA isolation of the epithelial monolayer was performed with the Qiagen Micro Kit. RNA library preparation and sequencing were performed (GeneWiz). Using poly dT priming (selects for mRNA during the reverse transcription to cDNA) and Illumina HiSeq PE125 sequencing approximately 20 million reads recovered per technical replicate. Raw reads were processed, and FPKM calculated for each gene via the NF-CORE rnaseq workflow (15).

### Statistical analysis

Linear Mixed-Random Effects modeling via the python statsmodels (16) package to determine the correlation between FPKM for a given gene and co-culture (respecting biological and technical replicates), with the following formula:

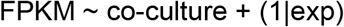

## Acknowledgements

Funding for these experiments, and *Bifidobacterium longum* (35624) were provided by Procter & Gamble; the design and execution of experiment as well as the analysis and interpretation of results were conducted independently of Procter & Gamble.

Dr. Golob is supported by the ASTCT New Investigator Award.

## References

1. Abrams GD, Bishop JE. 1967. Effect of the normal microbial flora on gastrointestinal motility. Proc Soc Exp Biol Med Soc Exp Biol Med N Y N 126:301–304.

2. Lomasney KW, Houston A, Shanahan F, Dinan TG, Cryan JF, Hyland NP. 2014. Selective influence of host microbiota on cAMP-mediated ion transport in mouse colon. Neurogastroenterol Motil Off J Eur Gastrointest Motil Soc 26:887–890.

3. Heneghan J. 2012. Germfree research: biological effect of gnotobiotic environments. Elsevier.

4. Mazmanian SK, Liu CH, Tzianabos AO, Kasper DL. 2005. An immunomodulatory molecule of symbiotic bacteria directs maturation of the host immune system. Cell 122:107–118.

5. Broom LJ, Kogut MH. 2018. The role of the gut microbiome in shaping the immune system of chickens. Vet Immunol Immunopathol 204:44–51.

6. Shimizu K, Muranaka Y, Fujimura R, Ishida H, Tazume S, Shimamura T. 1998. Normalization of reproductive function in germfree mice following bacterial contamination. Exp Anim 47:151–158.

7. Yan J, Charles JF. 2017. Gut Microbiome and Bone: to Build, Destroy, or Both? Curr Osteoporos Rep 15:376–384.

8. Warner BB. 2019. The contribution of the gut microbiome to neurodevelopment and neuropsychiatric disorders. Pediatr Res 85:216–224.

9. Hill DR, Spence JR. 2017. Gastrointestinal Organoids: Understanding the Molecular Basis of the Host-Microbe Interface. Cell Mol Gastroenterol Hepatol 3:138–149.

10. Fofanova T, Stewart C, Auchtung J, Wilson R, Britton R, Grande-Allen K, Estes M, Petrosino J. 2019. A novel human enteroid-anaerobe co-culture system to study microbial-host interaction under physiological hypoxia. preprint, Bioengineering.

11. Oeckinghaus A, Ghosh S. 2009. The NF-kappaB family of transcription factors and its regulation. Cold Spring Harb Perspect Biol 1:a000034.

12. Kawai T, Akira S. 2007. TLR signaling. Semin Immunol 19:24–32.

13. Shimamura S, Abe F, Ishibashi N, Miyakawa H, Yaeshima T, Araya T, Tomita M. 1992. Relationship between oxygen sensitivity and oxygen metabolism of Bifidobacterium species. J Dairy Sci 75:3296–3306.

14. Watson CL, Mahe MM, Múnera J, Howell JC, Sundaram N, Poling HM, Schweitzer JI, Vallance JE, Mayhew CN, Sun Y, Grabowski G, Finkbeiner SR, Spence JR, Shroyer NF, Wells JM, Helmrath MA. 2014. An in vivo model of human small intestine using pluripotent stem cells. Nat Med 20:1310–1314.

15. Ewels PA, Peltzer A, Fillinger S, Alneberg J, Patel H, Wilm A, Garcia MU, Di Tommaso P, Nahnsen S. 2019. *nf-core* : Community curated bioinformatics pipelines. preprint, Bioinformatics.

16. Seabold S, Perktold J. 2010. Statsmodels: Econometric and statistical modeling with python9th Python in Science Conference.

